# Celastrol alleviates SGLT2 inhibitor-induced diabetic hyperketonemia by inhibiting hepatic ketogenesis

**DOI:** 10.64898/2026.04.01.715734

**Authors:** Yinghan Zhu, Yiting Wang, Minglong Zhang, Lingxu Liu, Yang Tian, Zeyu Guo, Ran Zhang, Jinrui Zhang, Zhenyu Ma, Fude Fang, Li Yan, Xiaojun Liu

## Abstract

SGLT2 inhibitor (SGLT2i)-induced diabetic hyperketonemia is a life-threatening acute complication of diabetes. While Celastrol has been reported to exert beneficial effects on obesity; its potential role in ketogenesis remains unclear. In this study, Celastrol administration significantly attenuates the fasting-induced elevation of blood β-hydroxybutyrate. Moreover, a 7-day course of Celastrol (1 mg/kg/day) leads to reductions in body weight and fat mass. Mechanistically, Celastrol specifically downregulates HMGCS2 expression and suppressess hepatic ketogenesis through inhibiting PPARα expression in the short term (≤ 2 days). However, after prolonged treatment for 7 days, Celastrol modulates both PPARαand serum free fatty acids (FFAs) levels. Furthermore, anti-ketogenic effect of Celastrol is abolished in *Ppar*α□ */*□ mice. Importantly, Celastrol effectively ameliorates SGLT2i-induced hyperketonemia. In summary, Celastrol curbs hepatic ketone overproduction in a PPARα-dependent manner, indicating its protective potential against SGLT2i-induced hyperketonemia.

## Introduction

Ketogenesis primarily takes place in the hepatic mitochondrial matrix, producing soluble ketone bodies—acetone, acetoacetate (AcAc), and β-hydroxybutyrate (β-OHB)—which serve as energy substrates in extrahepatic tissues through the breakdown of free fatty acids (FFAs) [1, 2]. Under the catalysis of acyl-CoA synthetase, FFAs are activated to acyl-CoA, which then traverses the mitochondrial membrane and undergoes β-oxidation. The rate-limiting enzyme of ketogenesis, 3-hydroxy-3-methylglutaryl-CoA synthase 2 (HMGCS2), catalyzes the condensation of acetoacetyl-CoA (AcAc-CoA) and acetyl-CoA to form hydroxymethylglutaryl (HMG)-CoA. This intermediate is subsequently cleaved by hydroxymethylglutaryl-coenzyme A lyase (HMGCL) to release acetyl-CoA and AcAc. AcAc can be further reduced to β-OHB or spontaneously decarboxylated to acetone. The expression of HMGCS2 is transcriptionally activated during fasting through the action of peroxisome proliferator-activated receptor α (PPARα), a central hepatic regulator that orchestrates the adaptive response to starvation by modulating both β-oxidation and ketogenic pathways [3].

Ketogenesis is physiologically enhanced under conditions of depleted carbohydrate reserves or elevated fatty acid availability. However, excessive production of ketone bodies may result in ketoacidosis, a potentially fatal metabolic state characterized by the accumulation of acidic ketones. Diabetic ketoacidosis (DKA) represents an acute and life-threatening complication of diabetes, characterized by hyperglycemia, metabolic acidosis, and ketosis [4]. Euglycemic DKA (EDKA) shares the metabolic acidosis and ketosis of DKA but occurs in the absence of significant hyperglycemia (blood glucose <200 mg/dL) [5]. Because of its atypical presentation, EDKA is often undiagnosed or diagnosed late, delaying treatment. Known triggers of EDKA include reduced caloric intake, excessive alcohol consumption, chronic liver disease, glycogen storage disorders, and recent use of insulin or sodium-glucose cotransporter-2 inhibitors (SGLT2i) [5]. SGLT2i, a class of glucose-lowering agents used in type 2 diabetes (T2D), act by inhibiting renal glucose reabsorption. However, they are associated with an increased risk of DKA—including EDKA—mainly through promoting lipolysis, enhancing renal ketone reabsorption, and altering the glucagon/insulin ratio via stimulation of pancreatic α cells and suppression of β cells [5–7]. These risks highlight the clinical need for effective preventive and therapeutic strategies against SGLT2i-induced DKA.

Celastrol (Cel), a pentacyclic triterpenoid derived from the traditional Chinese medicine *Tripterygium wilfordii* Hook F., has garnered attention in recent years for its broad pharmacological effects across metabolic diseases and cancer [8, 9]. It has been identified as a leptin sensitizer and a potential treatment for obesity and hepatic steatosis [10–12]. The anti-obesity properties of Cel are attributed to its ability to regulate leptin sensitivity, energy metabolism, inflammatory responses, lipid metabolism, and gut microbiota composition [8]. Furthermore, Cel has also been shown to ameliorate insulin resistance and confer protection against T2D and its complications through multiple mechanisms including activation of the phosphoinositide 3-kinase (PI3K)/protein kinase B (AKT) signaling pathway, upregulation of glucose transporter 4 (GLUT4) expression, and suppression of nuclear factor kappa-B (NF-κB) activation [8, 13, 14]. However, its role in hepatic ketogenesis has remained largely unexplored.

In this study, we demonstrated that Cel attenuates fasting-induced upregulation of HMGCS2 in a PPARα-dependent manner, thereby effectively curbing hepatic ketone overproduction. Notably, Cel administration also ameliorated SGLT2i-induced ketoacidosis in a streptozotocin (STZ)-induced T2D mouse model.

## Materials and Methods

### Mice

Male C57BL/6 mice were purchased from Beijing HFK Bioscience. Co., Ltd. *Ppar*α*^-/-^*mice were obtained from Cyagen Biosciences. Mice (≤ 4/cage) were housed in the specific pathogen-free conditions facility and maintained on a 12 h light-dark cycle with free access to water and a normal chow diet (NCD, 9% fat; Lab Diet). Mice were treated with intraperitoneally Cel (Abmole, Cat#M3884) at the indicated concentration and time, followed by the specified fasting period. □ To generate an T2D mouse model, high-fat diet (HFD; 60% kcal fat, Research Diet)-fed mice were injected with low-dose streptozotocin (STZ). Briefly, 6-week-old male C57BL/6 mice were fed an HFD for 4 weeks, fasted for 12 hours, and then injected intraperitoneally with 40 mg/kg STZ (Abmole, Cat#M2082) for 5 consecutive days, and then maintained on HFD continuously for additional 2 weeks. For SGLT2i–induced hyperketonemia experiments, T2D mice pretreated with Cel (1 mg/kg/day) intraperitoneal for 7 days. Mice were subsequently injected intraperitoneally with dapagliflozin (1 mg/kg; Abmole, Cat# M1937) or vehicle (10% DMSO in PBS) after 4-hour food deprivation, followed by 8 hrs of fasting and water deprivation. Blood glucose and β-OHB levels were measured from tail blood using a One-Touch Ultra® glucometer (LifeScan Inc., Milpitas, CA) and FreeStyle Optium Neo system (Abbott), respectively. Serum FFAs concentrations were quantified using a commercial assay kit (Solarbio, BC0595-100T/96S). All animal experiments were conducted under protocols approved by the Animal Research Committee of the Institute of Laboratory Animals, Institute of Basic Medical Sciences Chinese Academy of Medical Sciences & School of Basic Medicine Peking Union Medical College (ACUC-A01-2022-010).

### Cell culture

Mouse primary hepatocytes (MPH) were isolated from 8-week-old male C57BL/6 mice or *Ppar*α*^-/-^* mice, and cultured in RPMI 1640 medium supplemented with 10% FBS as previously described [15]. After the MPH adhere to the surface, add the Cel to a final concentration of 10 μm or vehicle control (1% DMSO in PBS).

### Western blotting

Protein was extracted from frozen tissue samples in cell lysis buffer supplemented with proteinase inhibitor. The protein concentration of each lysate was determined using a BCA protein assay kit (LABLEAD, Cat#B5000) to ensure equal loading. Subsequently, the equal amount of protein was loaded onto a 10% SDS-polyacrylamide gel, and separated proteins were transferred to PVDF membranes. Western blot assays were performed using indicated specific antibodies including anti-HMGCS2 antibody (Abclonal, Cat#A14244), anti-PPARα antibody (Santa Cruz Biotechnology, Cat#sc-398394), anti-GAPDH antibody (CWBio, Cat#CW0100M). The proteins bands were quantified by ImageJ software.

### RNA sequencing (RNA-Seq)

The RNA-Seq was performed according to the manufacturer’s protocol (Annoroad Gene Technology Co.Itd). Briefly, total RNA was extracted from the liver of control and Cel (10 mg/kg) treatment mice (n=3/group) using Trizol for RNA-Seq to screen the differentially expressed genes (DEGs) (Invitrogen, Carlsbad, CA, USA) according to manual instruction. Total RNA was enriched by oligo (dT)-attached magnetic beads, followed by library construction and sequencing analysis. The data were mapped to mouse reference genome (GRCm39) by using Bowtie2. The data have been deposited to National Genomics Data Center, China National Center for Bioinformation (NGDC-CNCB) (https://ngdc.cncb.ac.cn/) with the dataset identifier CRA021250.

### Real-Time Quantitative PCR (RT-qPCR)

Total RNA was extracted from mouse tissues using a Trizol-based method. 2 μg of total RNA was reverse-transcribed into a first-strand cDNA pool using reverse transcriptase and random primers, according to the manufacturer’s instructions. RT-qPCR was performed using Hieff UNICON® Universal Blue qPCR SYBR Green Master Mix (Cat#11184ES03, Yeasen) with the gene-specific primers. All gene expression data were normalized to *Gapdh* expression levels. Primer sequences were as follows: *Hmgcs2*, GAGCGATGCAGGAAACTTCG and GTATCTGTTTTGGCCAGGGGA; *Hmgcl*, GTCTTCGGTGCTGTGTCTGA and TCTTGGCAACCTCAGCAACT; *Ppar*α, ACAAGGCCTCAGGGTACCA and GCCGAAAGAAGCCCTTACAG; *Gapdh*, TCTCCTGCGACTTCAACA and TGGTCCAGGGTTTCTTACT.

### Molecular docking of Celastrol and PPARα

To identify potential target proteins for Cel and explore its interaction with upstream proteins of PPARα, molecular docking analysis was conducted to predict Cel’s binding sites with its potential target proteins. The three-dimensional structure of Cel was obtained from PubChem (https://pubchem.ncbi.nlm.nih.gov). The three-dimensional structure (AF-Q07869-F1-v4) of the PPARα protein was obtained from the <u>AlphaFold Protein Structure Database</u>. Molecular docking simulations were carried out using CB-Dock2 (<u>CB-Dock2: An accurate protein-ligand blind docking tool</u>).

### Biotinylated-celastrol pull-down assay

The detailed procedure was performed according to a previous study [16]. Briefly, protein lysates of liver tissue isolated from wild-type C57 mice were incubated gently with either 10 μM Biotinylated-celastrol (Bio-Cel, synthesized by Xi’an Ruixi Biological Technology Co., Ltd.) or 10 μM biotin for 4 h, followed by slow incubation with the beads for 4 h. Before and after incubation with the Streptavidin Magnetic Beads, the beads were washed 10 min for three times quickly by vortexing in RIPA buffer (supplemented with 100× protease inhibitor cocktail and 100× PMSF). Beads were separated using a magnetic stand, and bound proteins were eluted by boiling in loading buffer. The eluates from each group were then analyzed by western blotting protocol.

### Statistical analysis

Data analyses were performed with Graphpad Prism 8. For data showing normal distribution, two-tailed unpaired Student’s *t* test was used to compare differences between two groups, and one-way ANOVA was used to compare differences between more than two groups. For data showing skewed distribution, a nonparametric statistical analysis was performed using the Mann-Whitney U test. All data were presented as the mean ± SD. * p < 0.05 was considered statistically significant.

## Results

### Celastrol suppresses fasting-induced hepatic ketogenesis

To investigate the effect of Cel on hepatic ketogenesis, male wild type (WT) mice were treated with either vehicle or Cel via intraperitoneal injection, and then were subjected to starvation (Figure 1A). We first assessed the effect of 10 mg/kg Cel on ketogenesis. A single intraperitoneal injection at 10 mg/kg, combined with concurrent 24-hour fasting, was found to reduce blood β-OHB levels (Figure 1B). We further examined the effects of administering 1 mg/kg/day and 3 mg/kg/day Celastrol over two days, along with a simultaneous 48-hour fasting. The results indicated that treatment with 3 mg/kg/day for two days—but not 1 mg/kg/day—attenuated the rise in blood β-OHB induced by the 48-hour fasting (Figure 1B). Finally, we investigated the longer-term impact of Celastrol at 1 mg/kg/day administered for 7 days, with fasting during the final 2 days. This extended treatment significantly suppressed the increase in β-OHB levels (Figure 1B). Additionally, administration of 1 mg/kg/day Cel for 7 days did not alter blood β-OHB levels under feeding condition (Supplementary Figures S1A,B). Consistent with previous report on its glucoregulatory properties [10], Cel treatment (3 mg/kg/day for 2 days under fasting condition, or 1 mg/kg/day for 7 days under fasting and feeding conditions) correlated with improved glycemic control, as evidenced by reduced blood glucose levels (Figure 1C and Supplementary Figure S1C). While short-term (≤ 2 days) Cel treatment did not alter body weight, long-term (7 days) treatment reduced it (Figure 1D and Supplementary Figure S1D). These findings establish Cel as a potent modulator of fasting-associated ketogenesis.

**Figure 1.**
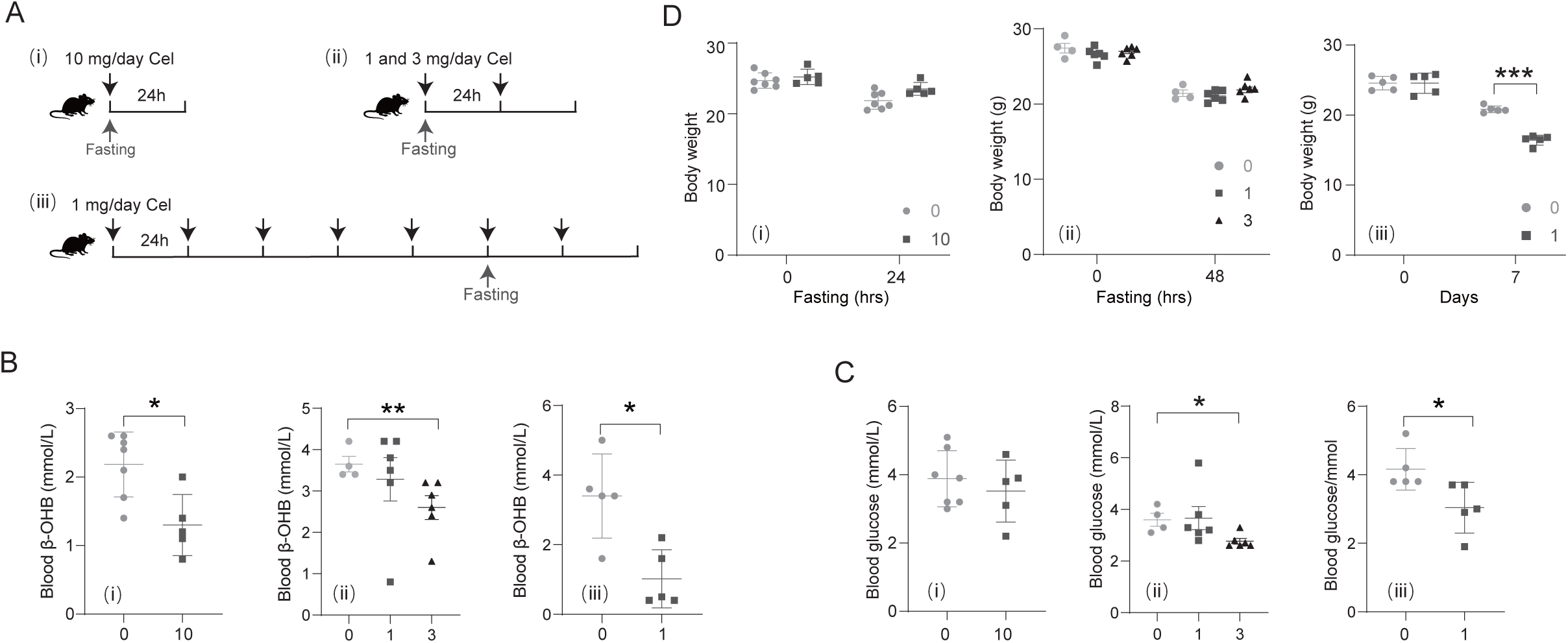
Celastrol suppresses fasting-induced hepatic ketogenesis. (A) Schematic of the experimental procedure about WT mice treated with Cel under three different regimens. (i) WT mice were treated with a single dose of Cel (10 mg/kg) combined with a 24-hour fast, Ctrl: n = 7, Cel: n = 5; (ii) WT mice were treated with Cel at doses of 1 or 3 mg/kg/day for 2 days combined with a 48-hour fasting, Ctrl: n = 4, 1 and 3 mg Cel: n = 6/group; (iii) WT mice were treated with Cel at a dose of 1 mg/kg/day for 7 days along with fasting during the last two days, n = 5/group. (B-D) Effects on blood β-OHB (B), blood glucose (C), and body weight (D) are shown. Data are presented as mean ± SD. *p < 0.05, **p < 0.01 and ***p < 0.001 by One-way ANOVA for D (ii), Student’s *t* test for B (i), B (iii), C (i), D (iii), and Mann Whitney test for B (ii), C (ii), C (iii) and D (i).

### Celastrol reduces the expression of HMGCS2

Given that liver is the principal site of systemic ketogenesis, we performed RNA-Seq analysis to identify DEGs following treatment with 10 mg/kg/day Cel, aiming to elucidate the mechanism by which Cel reduces hepatic ketogenesis. The data suggested that Cel administration resulted in 3922 DEGs (|logFC| ≥ 1, p < 0.05, q < 0.05), including 1691 upregulated and 2231 downregulated genes (Figure 2A and Supplementary Table S1). Gene Ontology (GO) enrichment analysis revealed significant alterations in metabolic pathways, particularly those involved in fatty acid metabolism, fatty acid beta-oxidation, cellular response to glucose and amino acid stimuli, and ketone body biosynthesis (Figure 2B,C and Supplementary Table S2). Among the key enzymes regulating hepatic ketogenesis, hepatic *Hmgcs2* and *Hmgcl* were downregulated, which were further confirmed by qPCR (Figure 2C-E). Furthermore, regardless of whether Cel was administered for short-term or long-term treatment, it reduced the mRNA levels of hepatic *Hmgcs2* and *Hmgcl* in mice under fasting or feeding conditions (Figures 2E and Supplementary Figure S2A). Additionally, Cel treatment increased the mRNA level of *Fabp4*, while decreasing the mRNA levels of *Slc27a5* and *Cd36*, which was also consistent with the results of RNA-seq (Supplementary Figure S2B). Cel also reduced hepatic HMGCS2 protein expression in the liver (Figure 2F). Similar suppression was observed in mouse primary hepatocytes (MPH) (Figures 2G,H). These results indicate that Cel inhibits ketogenesis by downregulating HMGCS2 and HMGCL expression.

**Figure 2.**
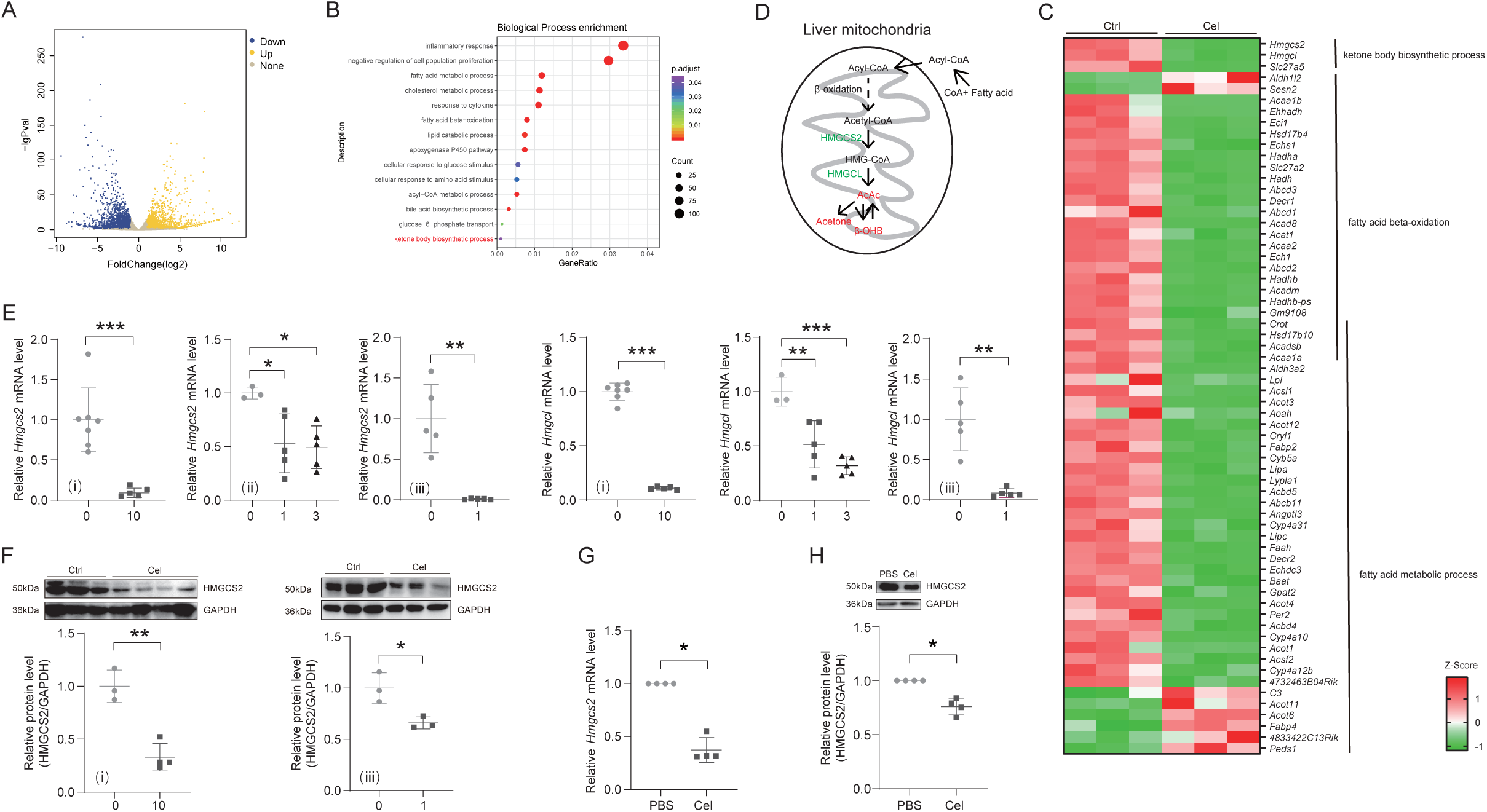
Celastrol reduces hepatic ketogenesis by inhibiting HMGCS2. (A-C) Volcano plot (A), GO enrichment analysis (B) and heatmap (C) of DEGs in the livers of mice treated with or without 10 mg/kg Cel, as indicated in Figure 1A (i). (D) Simplified schematic of ketone body production in liver. (E and F) Effects of different concentrations of Cel on the mRNA (E: (i) Ctrl: n = 7, Cel: n = 5; (ii) Ctrl: n = 3, 1 and 3 mg Cel: n = 5/group; (iii) n = 5/group) and protein (F: up, western blotting; down, quantitative result) levels of hepatic ketogenesis-related enzymes in mice treated as indicated in Figure 1A. (G and H) The mRNA (G, n =4/group) and protein (H: up, western blotting; down, quantitative result, 4 replicates) levels of HMGCS2 in MPH. Data are presented as mean ± SD. *p < 0.05, **p < 0.01 and ***p < 0.001 by One-way ANOVA for *Hmgcs2* and *Hmgcl* in E (ii), Student’s *t* test for E (i), E (iii) and F, and Mann Whitney for G and H.

### Celastrol inhibits the expression of PPARα

PPARα is a nuclear receptor that promotes ketogenesis through transcriptional activation of *Hmgcs2* [17]. In this study, Kyoto Encyclopedia of Genes and Genomes (KEGG) pathway analysis indicated Cel treatment significantly modulated the PPAR signaling pathway (Figure 3A,B and Supplementary Table S3). Both short- and long-term Cel treatment reduced the mRNA and protein levels of PPARα in the liver (Figures 3C,D). Moreover, Cel treatment for 7 days also decreased hepatic *Ppar*α mRNA level under feeding condition (Supplementary Figure S3A). Additionally, Cel treatment decreased the mRNA level of *Ppar*α in MPH (Figure 3E). As ligand-activated transcription factors, the activity of PPARs can be modulated by endogenous ligands and synthetic compounds [18]. Building upon previous finding that Cel binds to PPARγ and inhibits its transcriptional activity [19], we further investigated its potential multi-target pharmacological effects by performing molecular docking analysis between Cel and PPARα [20]. Molecular docking simulations revealed a high-affinity interaction between Cel and PPARα, with a Vina score of −8.2 kcal/mol (Figures 3F,G and Supplementary Figure S3B). The interaction between Cel and PPARα was further confirmed by biotinylated-Cel pull-down assay (Figure 3H).

**Figure 3.**
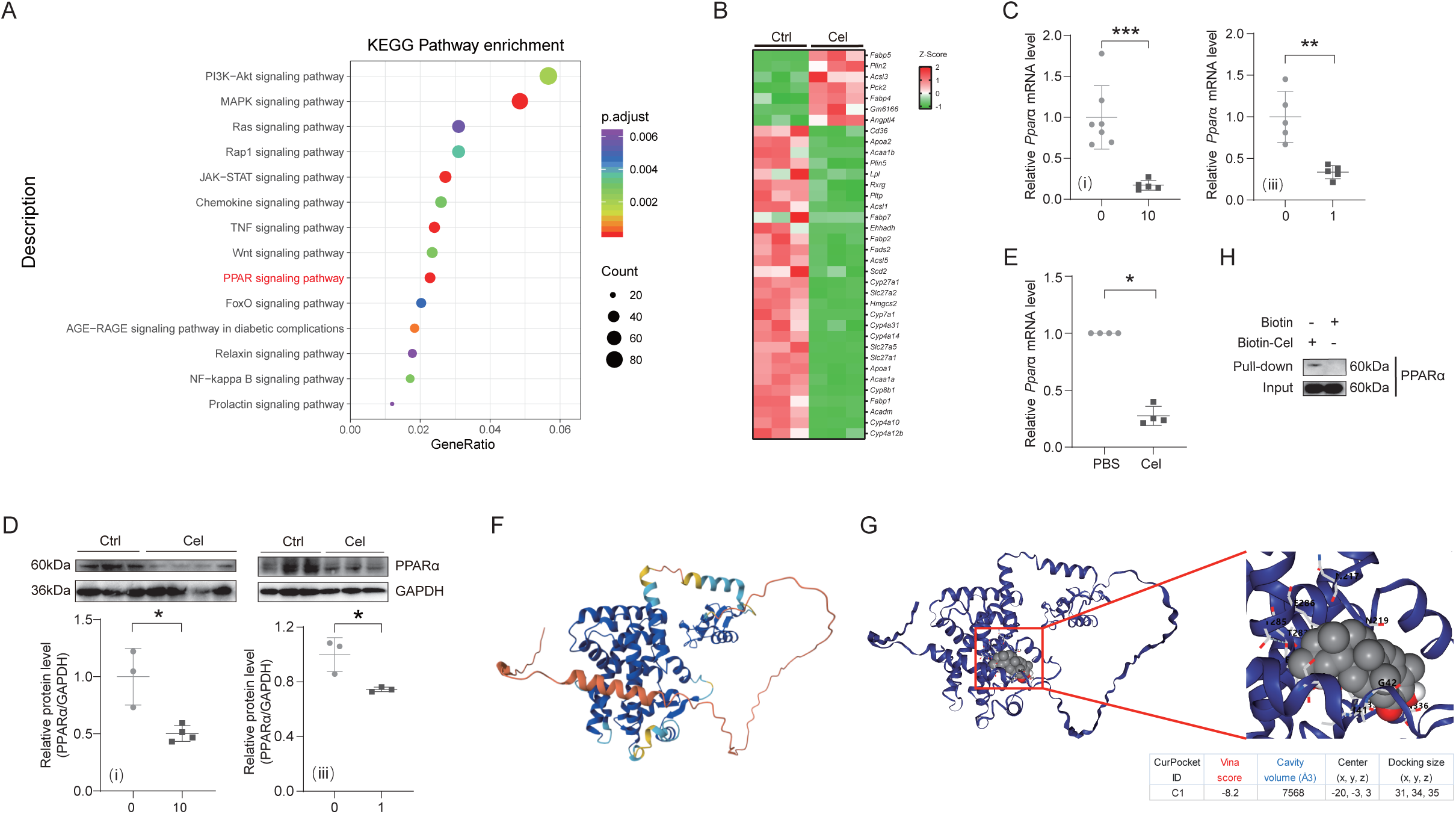
Celastrol inhibits the expression of PPARα. (A and B) KEGG pathway enrichment analysis (A) and heatmap (B) of DEGs from livers of mice treated with or without 10 mg/kg Cel as indicated in Figure 1A (i). (C and D) Different concentrations of Cel reduced the mRNA (C, (i) Ctrl: n = 7, Cel: n = 5; (iii) n = 5/group) and protein (D: up, western blotting; down, quantitative result) levels of hepatic PPARα in WT mice, as indicated Figure 1A (i) and (iii). (E) Cel reduced *Ppar*α mRNA levels in WT MPH. n =4/group. (F) Three-dimensional structure of the PPARα protein (PDB: AF-P23204-F1-v4). (G) The molecular docking structure of PPARα bound to Cel. (H) Physical association of PPARα and Cel confirmed by pull-down assay. Data are represented as mean ± SD. *p < 0.05 and ***p < 0.001 by Student’s *t* test for C and D and Mann Whitney test for E.

### Celastrol decreases hepatic ketogenesis dependent on PPARα

To further determine whether PPARα mediates the anti-ketogenic effect of Cel, *Ppar*α□ */*□ mice were used (Figure 4A). Genetic ablation of *Ppar*α completely abolished the ketogenesis-reducing effect of Cel, as evidenced by unaltered β-OHB levels and ketogenic enzyme expression in *Ppar*α□ */*□ mice (Figures 4B-D). Furthermore, Cel administration in *Ppar*α□ */*□ mice had no effect on blood glucose and body weight (Figures 4E,F). Additionally, Cel also did not affect HMGCS2 expression in *Ppar*α□ */*□ MPH (Figures 4G,H). Taken together, these results demonstrate that PPARα is essential for Cel-mediated suppression of hepatic ketogenesis.

**Figure 4.**
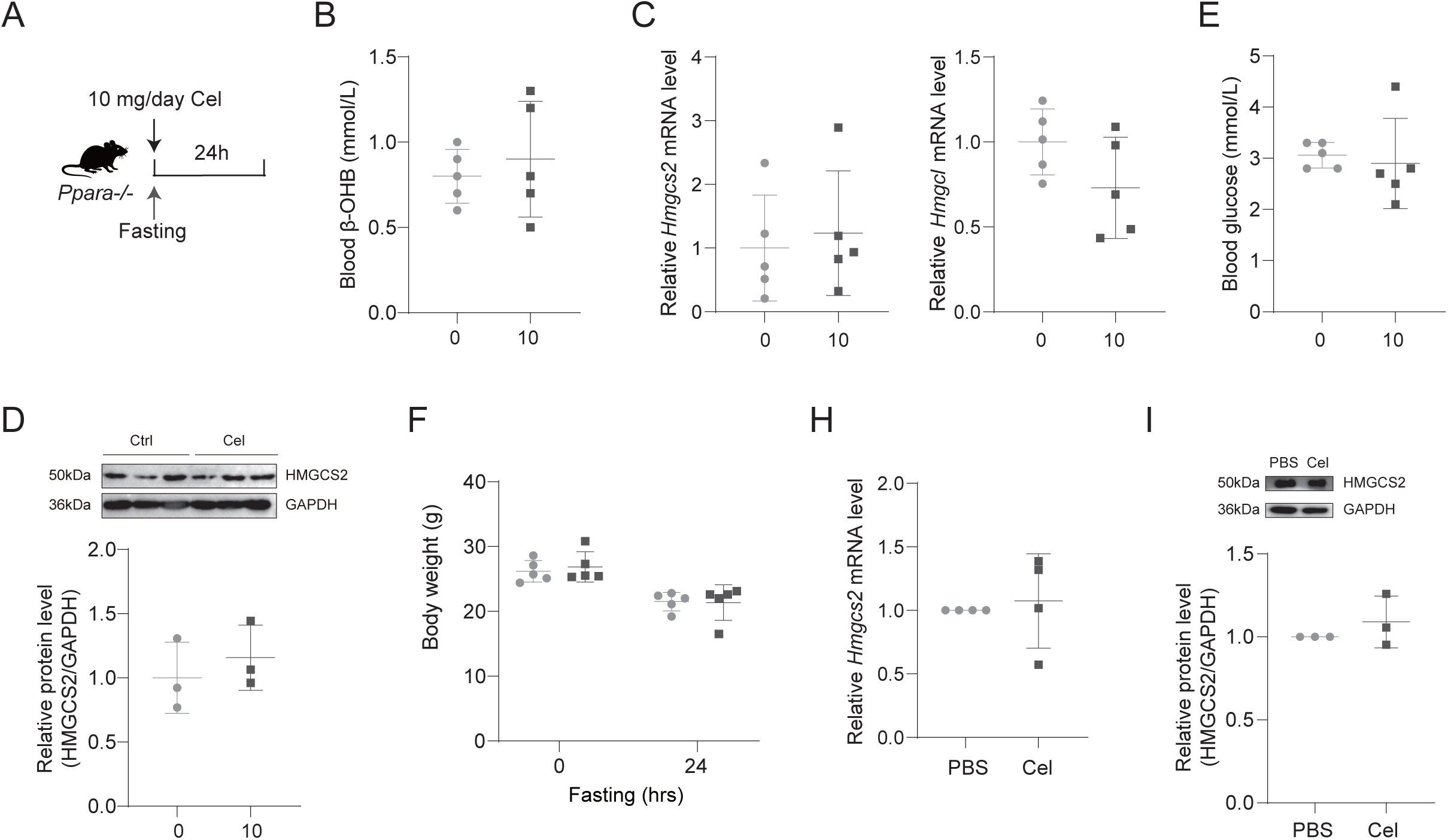
Celastrol decreases hepatic ketogenesis dependent on PPARα. (A) Schematic of the experimental procedure about *Ppar*α□ */*□ mice treated with or without a single dose of 10 mg/kg Cel combined with a 24-hour fasting. n = 5/group. (B-F) Cel treatment did not alter blood β-OHB (B), mRNA (C) and protein (D: up, western blotting; down, quantitative result) levels of hepatic ketogenesis-related enzymes, blood glucose (E), and body weight (F) in *Ppar*α□ */*□ mice treated as indicated in (A). (G and H) Cel treatment did not affect mRNA (G, n = 4/group) and protein (H: up, western blotting; down, quantitative result, 3 replicates) levels of HMGCS2 in *Ppar*α□ */*□ MPH. Data are represented as mean ± SD. The statistical analysis was performed by Mann Whitney test for G and H, and Student’s *t* test for the others.

### The short-term Celastrol treatment exerts hepatic anti-ketogenic effects independent of adipose tissue lipolysis

During fasting, adipose tissue lipolysis is activated, shifting whole-body fuel utilization from glucose and fatty acids to almost exclusively fatty acids after prolonged fasting [21]. These lipolysis-derived fatty acids are taken up by hepatocytes and converted into ketone bodies. Similar to the effect of Cel on body weight (Figure 1D), the short-term Cel treatment did not alter, whereas long-term treatment for 7 days reduced white adipose tissue (WAT) weight in WT mice under feeding and fasting conditions (Figure 5A and Supplementary Figure S4A). Furthermore, long-term Cel treatment for 7 days, but the short-term treatment, also decreased serum FFAs level in WT mice under feeding and fasting conditions (Figure 5B and Supplementary Figure S4B). These findings suggest that the short-term anti-ketogenic effect of Cel is primarily mediated through inhibition of hepatic ketogenic enzymes rather than suppression of adipose lipolysis.

**Figure 5.**
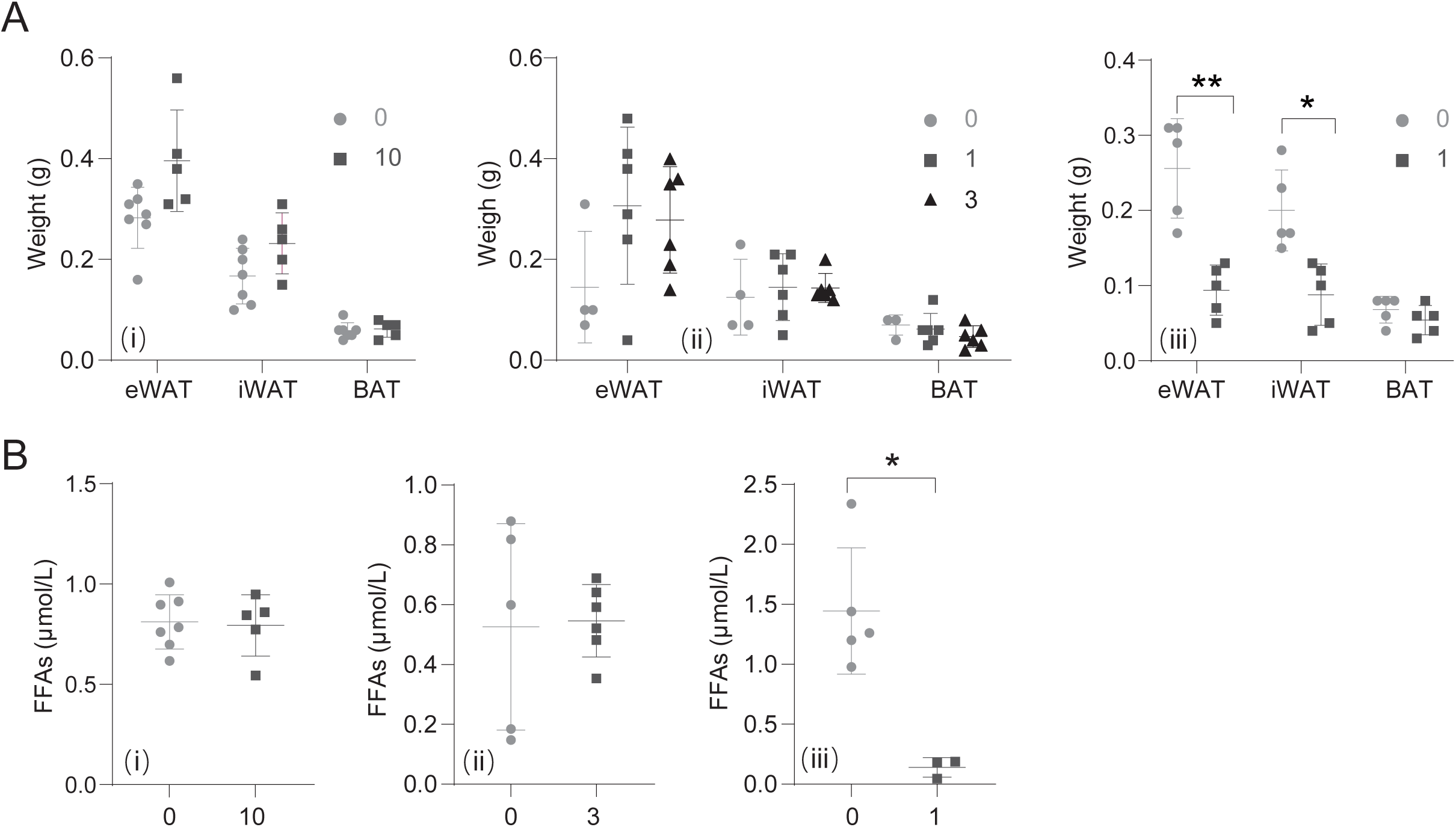
The effect of Celastrol treatment on adipose tissues weight and serum FFAs. (A and B) Effects of different concentrations of Cel on fat weights including epididymal white fat tissue (eWAT), inguinal white fat tissue (iWAT), and brown fat tissue (BAT) (A) and serum FFAs (B) in WT mice treated as indicated in Figure 1A. (ii) Ctrl: n = 7, Cel: n = 5; (ii) Ctrl: n = 4, 1 and 3 mg Cel: n = 6/group; (iii) n = 5/group. Data are represented as mean ± SD. * p < 0.05 and ** p < 0.01 by Mann Whitney test for A (i), A (ii) and B (iii), and Student’s *t* test for A (iii), B (i) and B (ii).

### Celastrol ameliorates SGLT2i-induced diabetic ketoacidosis

To address the clinical challenge of SGLT2i-induced hyperketonemia [22, 23], we adopted a pharmacological preconditioning strategy (Figure 6A). In STZ-induced diabetic mice, treatment with SGLT2i dapagliflozin alone did not significantly alter body weight and WAT weight, whereas pretreatment with Cel reduced both (Figures 6B,C). Consistent with a previous report [6], dapagliflozin treatment lowered blood glucose but elevated blood β-OHB and FFAs in STZ-induced T2D mice (Figures 6D-F). Preconditioning with Cel (1 mg/kg/day for 7 days) prior to dapagliflozin administration not only further lowered blood glucose but also attenuated dapagliflozin-induced ketogenesis (Figures 6D-F). Moreover, the increase in hepatic HMGCS2 and PPARα protein levels induced by dapagliflozin was reversed by Cel preconditioning (Figure 6G). Collectively, these results indicate that Cel may represent a promising adjunctive therapy to mitigate the risk of SGLT2i-associated hyperketonemia.

**Figure 6.**
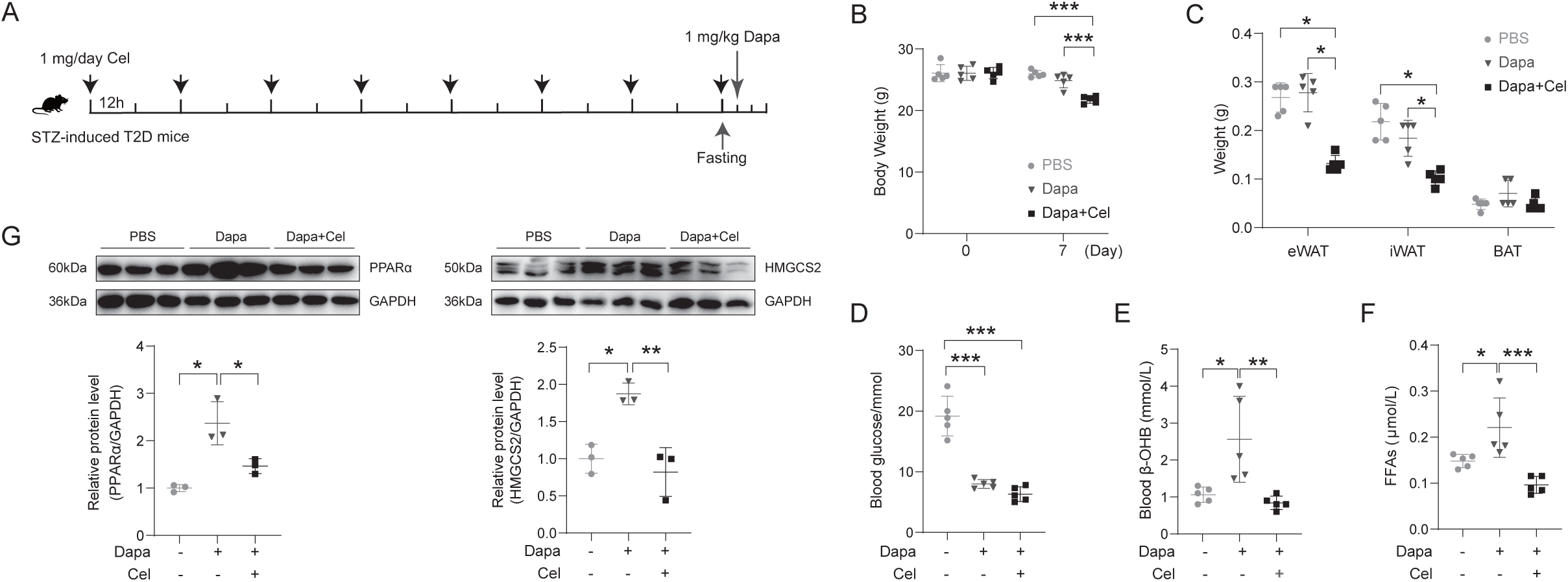
Celastrol ameliorates SGLT2i-induced diabetic ketoacidosis. (A) Schematic of the experimental procedure about STZ-induced T2D mice treated with Cel. The STZ-induced T2D mice were pretreated with or without Cel (1 mg/kg/day) for 7 days, followed by dapagliflozin administration after 4 hours of food deprivation and subsequent an 8-hour fasting period with water withdrawal. n = 5/group. (B-G) Body weight (B), fat tissues weight (C), blood glucose (D), blood β-OHB (E), FFAs levels (F), and the protein levels of hepatic HMGCS2 and PPARα (up, western blotting; down, quantitative result) (G) from the mice as indicated in (A). Data represent the mean ± SD. *p < 0.05, ** p < 0.01, and ***p < 0.001 by Mann Whitney test for C and One-way ANOVA for the others.

## Discussion

In this study, Cel pretreatment significantly attenuated the fasting-induced elevation of plasma ketone bodies while causing a modest reduction in plasma glucose concentrations. Through both in vivo and in vitro experiments, we demonstrated that Cel suppresses hepatic ketogenesis in a PPARα-dependent manner, as well as by reducing serum FFAs. Consequently, Cel treatment ameliorated SGLT2i-induced hyperketonemia.

The regulation of ketogenesis involves four hierarchical mechanisms: (i) hormonal control, (ii) transcriptional regulation, (iii) post-translational modifications of enzymes, and (iv) biochemical substrate dynamics [17]. While multiple hormones including glucagon, cortisol, thyroid hormones, and catecholamines can modulate ketogenesis, insulin serves as the principal regulator. Transcriptional control of ketogenic enzymes constitutes another essential layer of regulation, with HMGCS2 acting as the rate-limiting enzyme. The expression of *Hmgcs2* is modulated by transcription factors such as PPARα, cyclic-AMP response binding protein (CREB), transcription specificity protein 1 (SP1), forkhead box A2 (FOXA2), and hepatocyte nuclear factor 4α (HNF4α) [17]. Notably, the promoter region of *Hmgcs2* contains peroxisome proliferator response elements (PPREs), allowing PPARα/RXR heterodimers to initiate transcription upon ligand binding. Interestingly, HMGCS2 also contributes to an autoregulatory feedback loop by forming a nuclear complex with PPARα, thereby enhancing its own transcription via PPRE binding [24]. Additionally, a recent study also demonstrated that PPARγ can bind to the promoter region of *Hmgcs2* and induce its expression [25]. The PPAR family including PPARα, PPARβ/δ, and PPARγ are ligand-activated transcription factors belonging to the nuclear receptor superfamily. Although all the PPARs are expressed across various tissues, PPARα expression is highest in the liver and brown adipose tissue [18]. PPARs regulate numerous cellular and metabolic processes by modulating the transcription of genes involved in lipid and glucose metabolism. Consequently, they represent important molecular targets for developing therapies against metabolic disorders such as type 2 diabetes, dyslipidemia, atherosclerosis, and metabolic syndrome [18, 26, 27]. While the in vivo relevance of PPAR activation by certain compounds remains unclear, PPAR activity can be modulated by endogenous ligands including fatty acids and fatty acid-derived compounds, as well as by synthetic compounds referred to peroxisome proliferators such as phthalates, and hypolipidemic fibrate drugs [18]. A previous work demonstrated that Cel binds to PPARγ and inhibits its transcriptional activity in HepG2 and 3T3-L1 cells [19]. Here, our results reveal a high-affinity interaction between Cel and PPARα, extending this understanding that Cel may also interact with PPARα and suppresses its transcriptional function.

Fasting markedly activates adipose tissue lipolysis, leading to elevated plasma FFAs and enhanced hepatic ketogenesis. PPARα is a master regulator of hepatic nutrient metabolism during fasting, promoting fatty acid oxidation and ketogenesis while modulating glucose production [18, 28]. Accordingly, fasted *Ppar*α-/- mice display a host of metabolic abnormalities such as hypoglycemia, hypoketonemia, and elevated plasma FFAs [28]. As the synthetic PPARα agonists, fibrates exert lipid-lowering effect by activating PPARα. Notably, fibrates cause hepatomegaly and peroxisome proliferation in mice, but raise plasma high density lipoprotein and reduce plasma triglycerides in humans, suggesting its therapeutic value in the treatment of dyslipidemia ^[18, 29]^. Our findings indicate that short-term Cel treatment selectively suppresses hepatic ketogenesis through PPARα inhibition, without initially affecting adipose lipolysis. Prolonged administration, however, modulates both PPARα signaling and adipose lipolysis dynamics. Although Cel and fibrates have opposite effects on the regulation of PPARα, both exhibit lipid-lowering properties. The occurrence of this phenomenon might be related to the multiple roles of Cel in regulating lipid metabolism. Collective evidence has positioned Cel as a compound with considerable potential for treating a range of metabolic disorders [8]. In this study, 7-day Cel treatment decreased both WAT weight and serum FFAs. The study from Liu et al. reported that Cel potentiates leptin sensitivity and mitigates obesity by activating the leptin receptor-signal transducer and activator of transcription 3 (STAT3) pathway and reducing hypothalamic endoplasmic reticulum stress [10]. Ma et al. demonstrated that Cel promotes energy expenditure, induces inguinal WAT browning and brown adipose tissue (BAT) activation, and upregulates fatty acid metabolism and mitochondria-related genes transcription via activating heat shock factor 1 (HSF1)-peroxisome proliferator-activated receptor γ coactivator-1α (PGC1α) transcriptional axis [11]. Our earlier work also showed that Cel attenuates hepatic lipogenesis and serum FFAs through silent mating type information regulation 2 homolog 1 (SIRT1), ameliorating liver metabolic damage [12]. Notably, Chellappa et al. demonstrated that Cel suppresses food intake selectively in aged mice, but not in young mice [30]. Additionally, Choi et al. found that Cel inhibits adipogenesis and increases FFAs in 3T3-L1 adipocytes by suppressing PPARγ2- and/or CCAAT/enhancer-binding protein α (C/EBPα)-induced transcriptional activity [31]. Collectively, these studies highlight multiple pathways through which Cel modulates lipid metabolism.

Cel exhibits potential pharmacological activities in a variety of diseases, including cancer, obesity, diabetes, and rheumatic and immune diseases [32, 33]. In this study, we evaluated the effects of the different doses Cel on ketogenesis. The results showed that treatments such as a single administering of 10 mg/kg Cel combined with concurrent 24-hour fasting, 3 mg/kg/day Cel over two days along with simultaneous 48-hour fasting, and 1 mg/kg/day Cel for 7 days with fasting during the final 2 days, all reduced blood β-OHB levels. Moreover, preconditioning with 1mg/kg Cel also ameliorated SGLT2i-induced hyperketonemia. Numerous studies have demonstrated that long-term administration of Cel (generally more than 3 weeks), either orally (1 ∼ 10 mg/kg/day) or via intraperitoneal injection (0.1 ∼ 0.5 mg/kg/day), improves the metabolic phenotype of mice and rats [8]. Additionally, a single intraperitoneal dose of 4.5 mg/kg administered exerted significant neuroprotective effects in a cerebral ischemia-reperfusion injury mice model [34]. However, the application of Cel is largely limited by its toxic side effects, including cytotoxicity, hepatotoxicity, and even neurotoxicity at high concentrations or with prolonged exposure [32]. After prolonged fasting more than one day, activation of adipose tissue lipolysis shifts whole-body fuel utilization from a mix of glucose and fatty acids in the fed state toward a nearly exclusive reliance on fatty acids [21]. To meet energy demands under these conditions, the liver supplies glucose and ketone bodies to peripheral organs by activating glycogenolysis, gluconeogenesis, β-oxidation, and ketogenesis [6]. Consistent with prior reports [35–37], Cel lowered blood glucose levels in our study. Moreover, Cel has been shown to enhance glucose uptake in hepatocytes, adipocytes, and myotubes via activation of the phosphatidylinositol-3-kinase (PI3K)-AKT signaling pathway and glucose transporter 4 (GLUT4) pathway, thereby improving insulin resistance [8]. Given that Cel reduces both circulating glucose and ketone bodies, its use raises potential safety considerations—particularly the risk of inadequate energy supply to the brain, which depends on glucose and ketones as primary fuel sources [18]. Several strategies, including combination therapy, novel dosage forms, and innovative drug delivery routes, have been employed to mitigate the toxicity and enhance the efficacy of Cel [32]. Metabolic dysfunction-associated steatotic liver disease (MASLD) arises from increased lipid deposition driven by diet, de novo lipogenesis (DNL), and FFAs mobilization from adipose tissue lipolysis [38]. Under sufficient hepatic ketogenesis conditions, FFAs are efficiently channeled into the ketogenesis pathway, thereby protecting the liver from excessive lipid accumulation. Conversely, insufficient hepatic ketogenesis promotes lipid accumulation through increasing DNL and inducing mitochondrial stress and dysfunction [39]. Thus, the liver lipotoxicity and MASL progression caused by impaired hepatic ketogenesis due to long-term Cel administration warrants careful consideration.

Dapagliflozin, a highly potent and reversible SGLT2 inhibitor, is globally approved for the treatment of T2D. Although rare, a low risk of DKA was more common with dapagliflozin than placebo [40]. A previous study showed that dapagliflozin could retard atherosclerotic progression by suppressing PPARα signaling pathway [41]. In contrast, a recent study reported that dapagliflozin attenuates lipotoxic tenocyte injury through upregulation PPARα [42]. Similar to this recent report, our study also demonstrates that dapagliflozin enhanced PPARα expression. Furthermore, several studies have shown that dapagliflozin can induce both the expression and activity of PPARγ [43–47]. Importantly, our results indicate that Cel treatment counteracts the dapagliflozin-induced increase in PPARα expression. In summary, this study demonstrates that Cel suppresses hepatic ketogenesis in a PPARα-dependent manner and identifies Cel as a promising therapeutic candidate for the prevention of SGLT2i-associated hyperketonemia (Figure 7).

**Figure 7.**
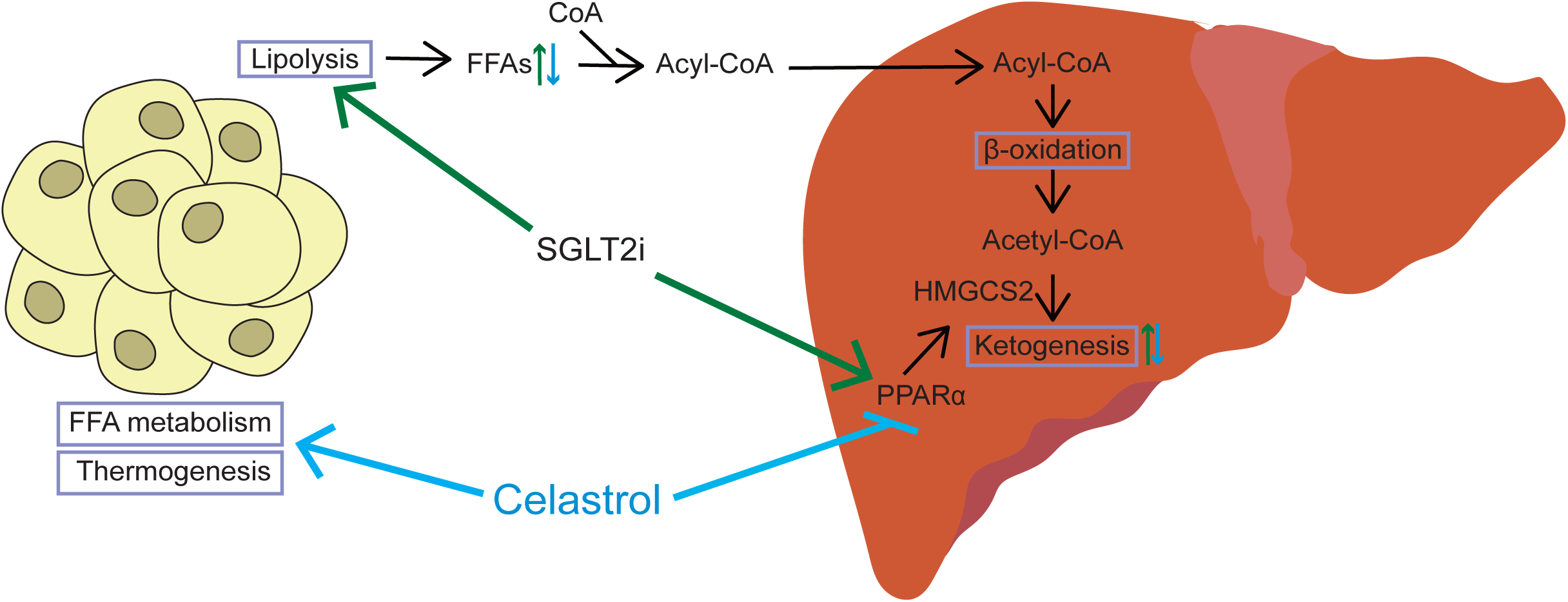
Proposed model illustrating how Cel regulates hepatic ketogenesis.

## Limitation of the study

Several limitations merit discussion. Firstly, although we identified the interaction with PPARα as a key mechanism, the detailed structural characterization of Cel-PPARα binding interface remains to be elucidated, future work about validation the binding site through mutagenesis experiments should be further performed. Second, while Cel was shown to reduce the mRNA level of *Ppar*α, the precise mechanism by which it regulates *Ppar*α gene transcription requires further clarification. Finally, although Cel has exhibited the promising efficacy in regulating glycolipid metabolism in mice, its pharmacokinetic properties and long-term safety profile warrant additional preclinical and clinical evaluation.

## Supplementary Data

Supplementary Data is available at Acta Biochimica et Biphysica Sinica online.

## Acknowledgments

We thank State Key Laboratory of Common Mechanism Research of Major Diseases Platform for consultation and instrument availability that supported this work. We thank Anoroad gene technology (Beijing) for the collaborative efforts in RNA-Seq.

## Author Contributions

Investigation and Methodology, Y.Z., Y.W., M.Z., L.L., Y.T., Z.G., R.Z, J.Z. and Z.M.; Writing - original draft, Formal analysis and Data curation, Y.Z.; Conceptualization, F.F.; Writing - review & editing, Supervision, Project administration and Funding acquisition, L.Y. and X.L. All authors reviewed the manuscript.

