## Supplementary material for "Celastrol alleviates SGLT2 inhibitor-induced diabetic hyperketonemia by inhibiting hepatic ketogenesis": Supplementary data.docx

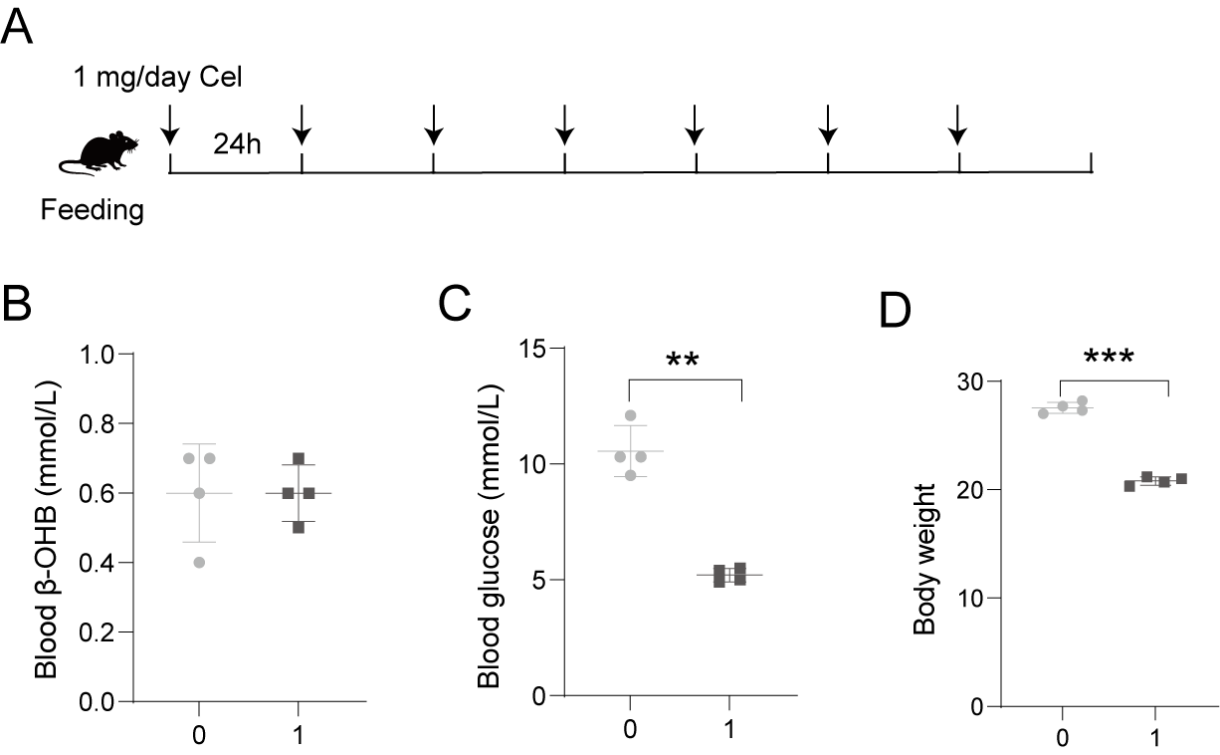

**Supplementary Figure S1. The effects of Celastrol on wild type mice under feeding condition.** (A) Schematic of the experimental procedure about WT mice treated with Cel at 1 mg/kg/day for 7 days. (B-D) Effects on blood β-OHB (B), blood glucose (C), and body weight (D) are shown. Data are presented as mean ± SD. n = 4/group. **p < 0.01 and ***p < 0.001 by Student’s *t* test.

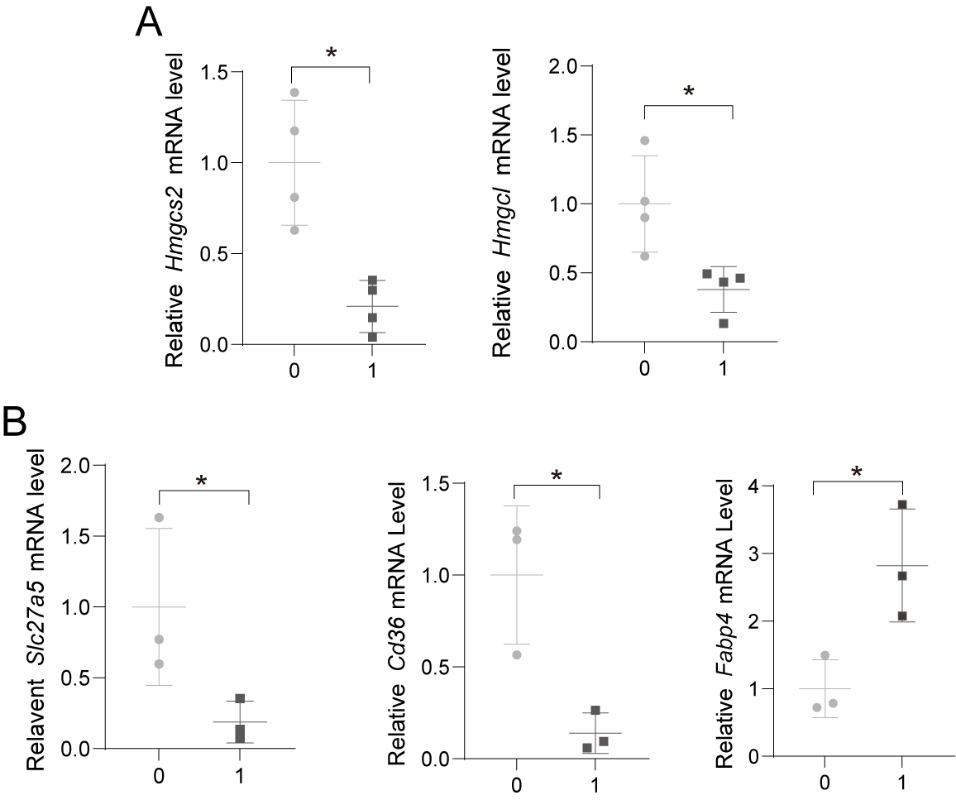

**Supplementary Figure S2. Celastrol regulates the mRNA levels of genes involved in hepatic ketogenesis and fatty acid metabolism.** (A) Cel reduces the mRNA levels of hepatic *Hmgcs2* and *Hmgcl*. WT mice were treated with Cel at a dose of 1 mg/kg/day for 7 days under feeding conditions. n =4/group. (B) Cel regulates the mRNA levels of hepatic *Slc27a5*, *Cd36* and *Fabp4*. WT mice were treated with Cel at a dose of 1 mg/kg/day for 7 days along with fasting during the last two days. n =3/group. Data are represented as mean ± SD. *p < 0.05 by Mann Whitney test for *Hmgcl* and by Student’s *t* test for the others.

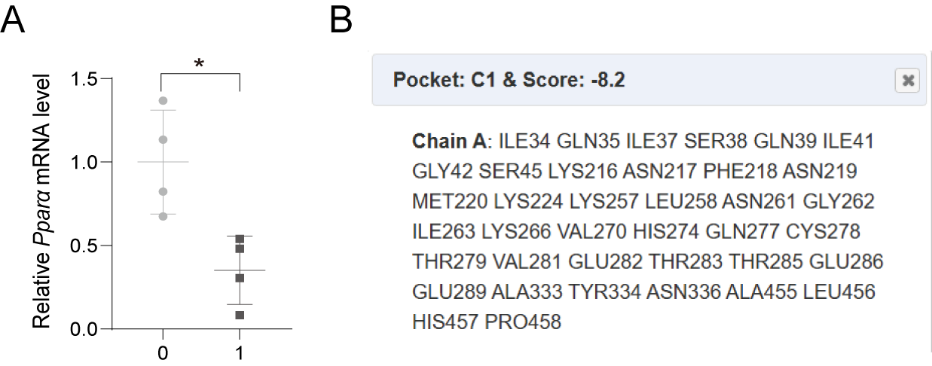

**Supplementary Figure S3. Celastrol inhibits the expression of PPARα.** (A) Celastrol inhibits the mRNA level of hepatic *Pparα* under feeding condition. 1 mg/kg/day Cel treatment for 7 days reduced the mRNA level of hepatic *Pparα* in WT mice. Data are represented as mean ± SD. *p < 0.05 by Student’s *t* test. (B) Cel contacts residues of PPARα by molecular docking.

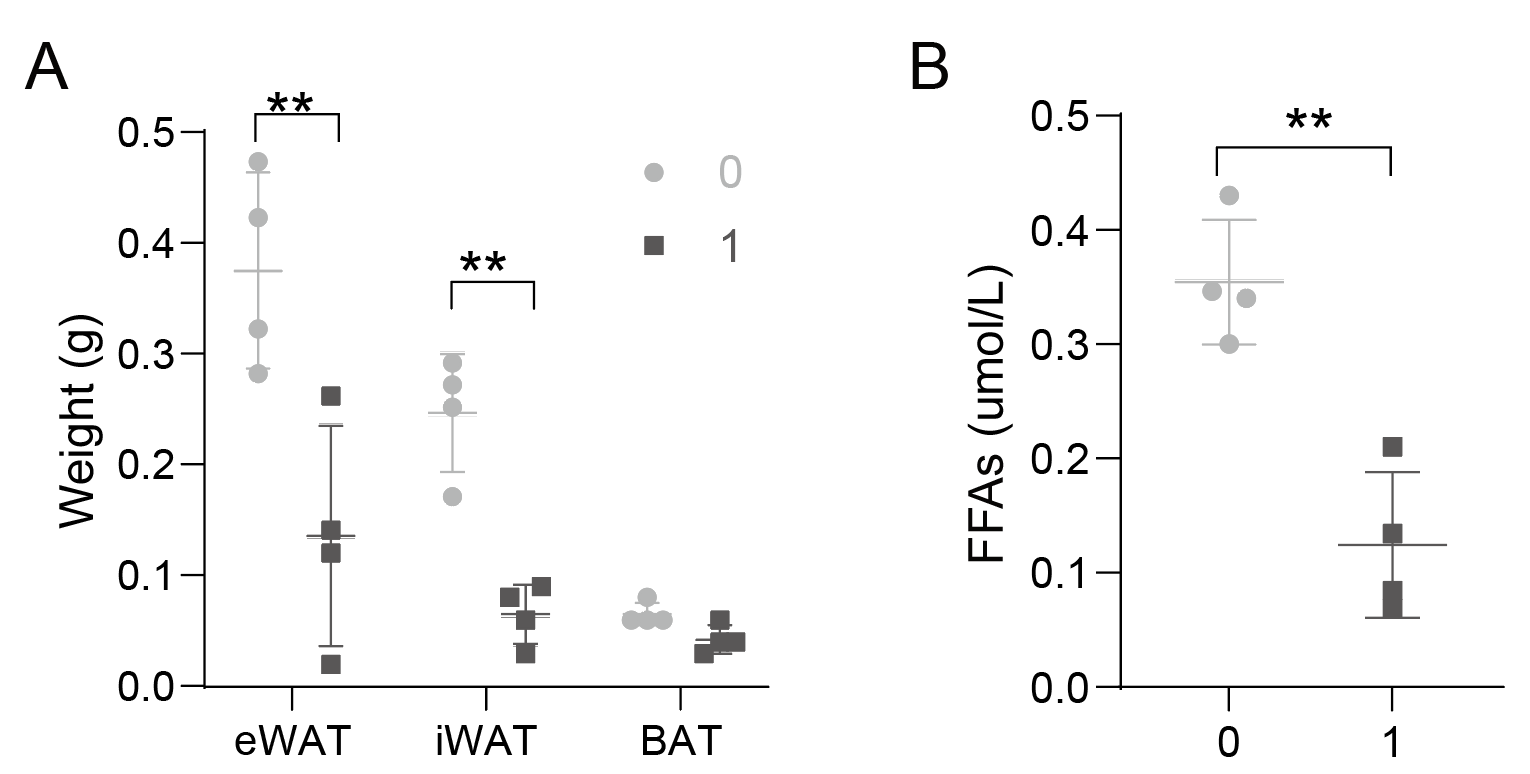

**Supplementary Figure S4. The effect of Celastrol treatment on adipose tissues weight and FFAs.** Treatment with Cel at 1 mg/kg/day for 7 days reduced fat weights (A) and serum FFAs (B) in WT mice under feeding conditions. Data are represented as mean ± SD. ** p < 0.01 by Student’s *t* test.
